## Supplementary Information for "Proteome allocation is linked to transcriptional regulation through a modularized transcriptome"

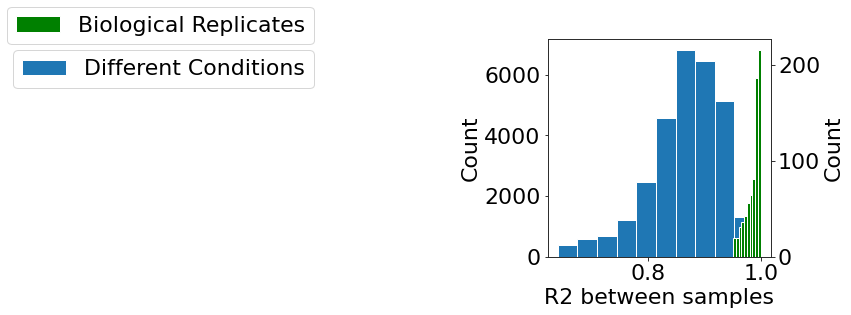

**Supplementary Figure 1. PRECISE1k replicate correlations.** R^2^ values between biological replicates and random samples within PRECISE1k.

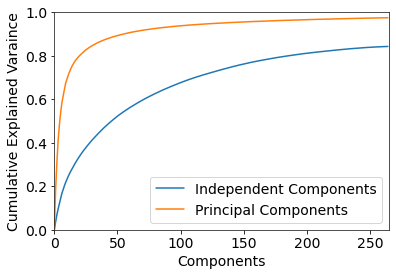

**Supplementary Figure 2. PRECISE1k explained variance.** Explained variance plot for PRECISE1k and its components.

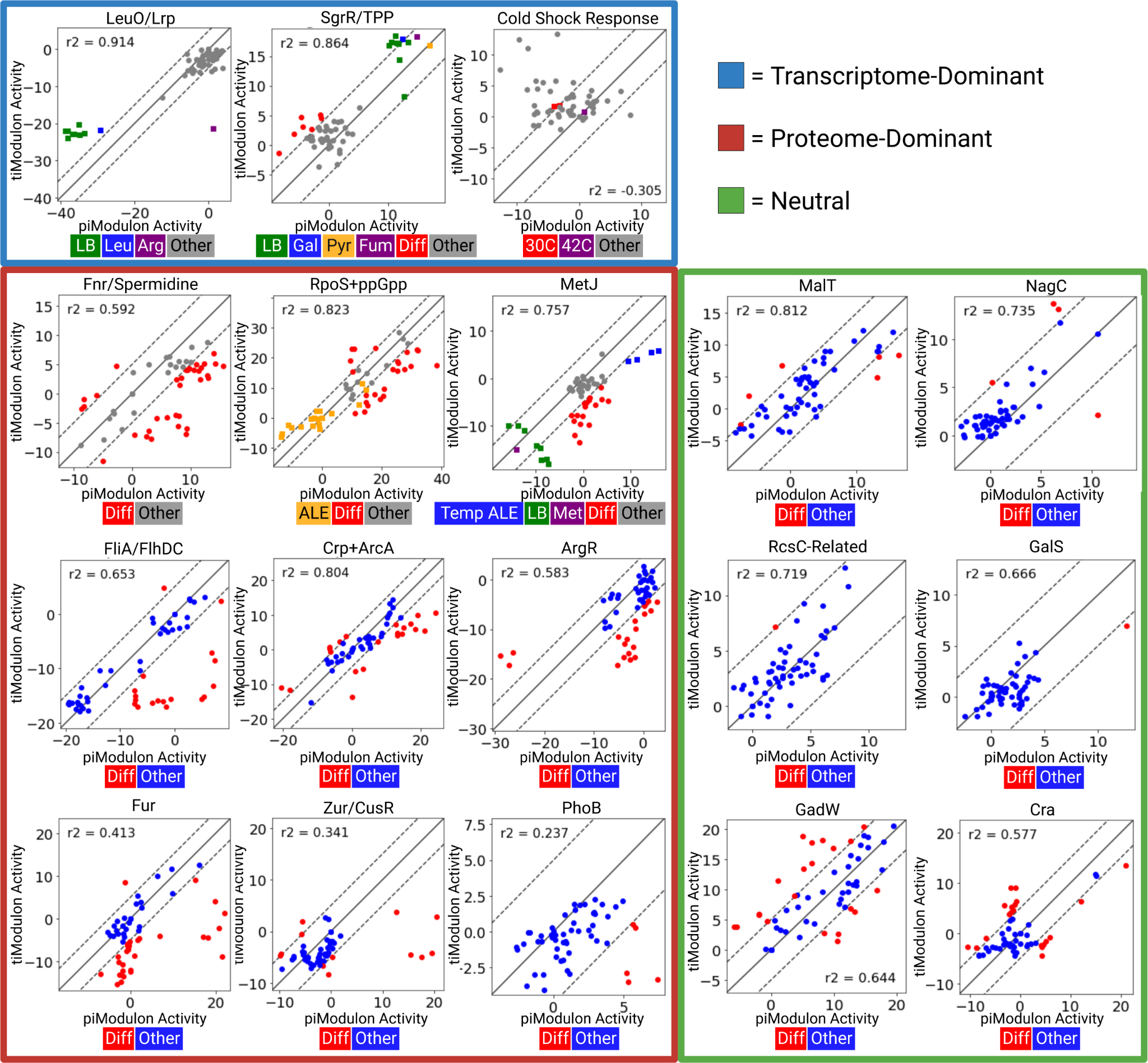

**Supplementary Figure 3. Comparing piModulon and tiModulon activities for all matched pairs.** Differential iModulon Activity (DiMA) plots for all matched iModulons between the two datasets. iModulons that are transcriptome-dominant (signal more active in the tiModulon than the piModulon) are highlighted in blue. iModulons that are proteome-dominant are highlighted in red, and iModulons that are neutral are highlighted in green. Activities are considered differentially activated for samples that lie outside the significance threshold (dashed line). Legends for each plot are placed below each plot. *Abbreviations LB: Lysogeny Broth, Leu: Leucine Supplement, Arg: Arginine Supplement, Gal: Galactose Carbon Source, Pyr: Pyruvate Carbon Source, Fum: Fumerate Carbon Source, Diff: Differentially Activated, Temp: Temperature, Met: Methionine Supplement.*

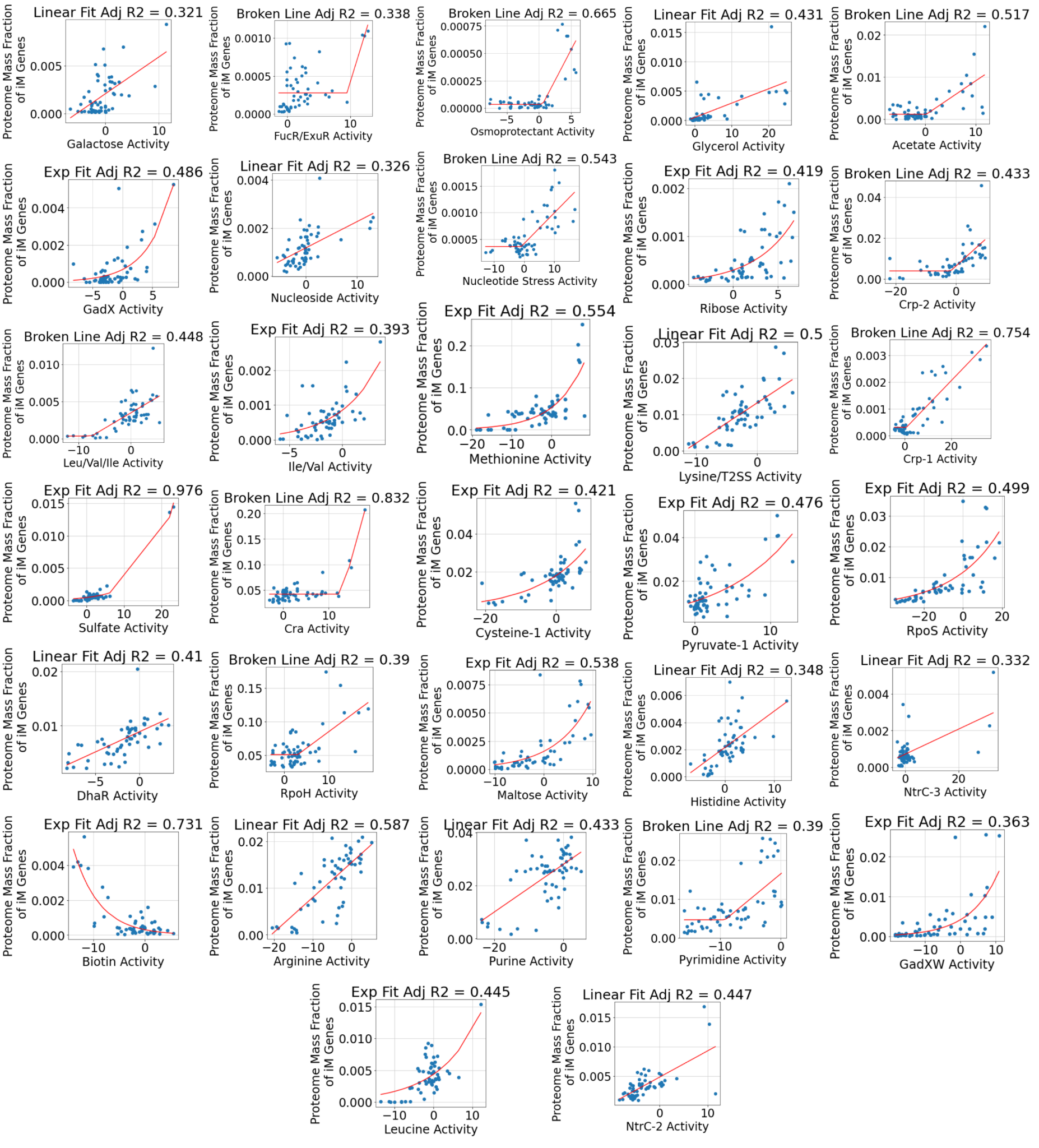

**Supplementary Figure 4. Scatter plots and regressions for all tiModulons with strong correlations.** Scatter plots for all tiModulon activities and their measured proteome mass fraction of the associated genes for tiModulons with strong correlations. tiModulons are charaterized based on which regression method (linear, exponential, and broken line) resulted in the best adjusted R^2^ value. The regression method and R^2^ value can be found above each plot, with the tiModulon name below. Individual samples are scatter plotted in blue, and the regression line is in red.

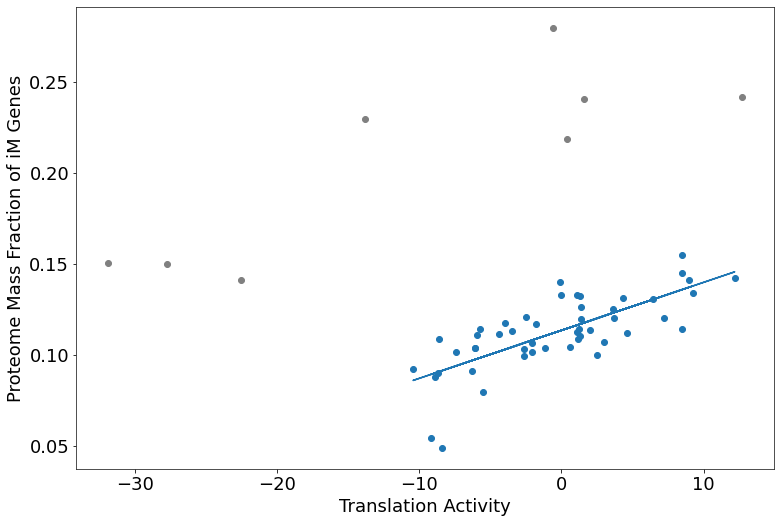

**Supplementary Figure 5. Proteome allocation of the translation tiModulon.** PRECISE1k Translation tiModulon activities plotted against their associated proteome mass fractions. There is a strong correlation between samples except for a few outliers (gray). R^2^ = 0.737 without outliers, R^2^ = 0.135 with outliers.

**Supplementary Table 1. Matched tiModulons and piModulons**

| **piModulon** | **tiModulon** | **Correlation** |
| --- | --- | --- |
| MalT | Maltose | 0.517757 |
| RcsC-related | UC-4 | 0.503688 |
| RcsC-related | pts ALE | 0.505555 |
| RpoS+ppGpp | RpoS | 0.63325 |
| NagC | GlcNAc | 0.656724 |
| FliA/FlhDC | FlhDC-2 | 0.336105 |
| FliA/FlhDC | FliA | 0.792489 |
| Fur | Fur-1 | 0.68668 |
| GalS | Galactose | 0.535344 |
| ArgR | Arginine | 0.332504 |
| Cold Shock Response | Cold Shock | 0.489819 |
| SgrR/Thiamine diphosphate | Thiamine-2 | 0.260406 |
| SgrR/Thiamine diphosphate | Thiamine-1 | 0.65899 |
| PhoB | Phosphate-1 | 0.427802 |
| PhoB | tpiA KO | 0.332344 |
| Zur/CusR | Zinc-1 | 0.411696 |
| Zur/CusR | Copper | 0.2603 |
| GadW | GadXW | 0.581244 |
| Translation-Null | Translation | 0.391143 |
| Fnr/Spermidine | Fnr-1 | 0.347905 |
| Fnr/Spermidine | Fnr-3 | 0.401703 |
| LeuO/Lrp | Lrp | 0.65983 |
| LeuO/Lrp | Pyruvate-1 | 0.264165 |
| Crp+ArcA | Crp-2 | 0.532799 |
| Crp+ArcA | Maltose | 0.255092 |
| Cra | Cra | 0.680723 |
| MetJ | Methionine | 0.448365 |
| Fimbriae | Fimbriae | 0.471338 |
